# MSACLR: Contrastive Learning of Protein Conformations from MSAs

**DOI:** 10.1101/2025.10.01.679854

**Authors:** Junjie Zhang, Enming Xing, Xiaolin Cheng

## Abstract

We propose **MSACLR** (**M**ultiple **S**equence **A**lignment **C**ontrastive **L**earning **R**epresentation), a two-stage contrastive learning framework that maps MSA space to conformational space. In Stage 1, embeddings are trained to discriminate structural folds across diverse proteins using only MSA information. In Stage 2, embeddings are fine-tuned on subMSAs labeled by their associated predicted structural clusters, enabling discrimination of alternative conformations within the same protein. To enrich training data, we introduce BLOSUM62-guided [1] augmentation, which expands the pool of subMSAs associated with each structural cluster label by introducing sequence-level diversity. Our experiments show that MSACLR embeddings achieve clearer fold-level separation than single-sequence baselines, while fine-tuned embeddings capture conformational variation across scales—from local loop motions to domain motions and fold switching. MSACLR provides a foundation for efficient exploration of MSA space and enables sampling of conformational ensembles, bridging the gap between static structure prediction and dynamic protein behavior.

## 1 Introduction

Protein structures are fundamental to biological function, and recent advances in deep learning - most notably AlphaFold2 [2] - have transformed the accuracy of protein structure prediction. Despite this breakthrough, current models primarily generate static snapshots, whereas proteins in their native environments exist as dynamic ensembles. Capturing alternative conformations is therefore essential for understanding molecular mechanisms and guiding therapeutic design, yet reliable multi-conformation prediction remains an open challenge.

Several strategies have recently been proposed to sample protein conformations beyond the static predictions of AlphaFold2. The first direction integrates AI predictions with enhanced sampling, as in AF2RAVE [3], which seeds simulations with MSA-perturbation ensembles to refine conformational landscapes. A second direction adapts structure prediction models directly, as exemplified by approaches such as CFold [4], which retrain AlphaFold2 on dataset containing multiple conformational states. A third direction repurpose folding as generative problem, with methods such as Alphaflow [5] and BioEmu [6] employing flow-matching or diffusion frameworks to generate diverse conformations conditioned on sequence inputs. A fourth direction focuses on perturbing MSA inputs to AlphaFold2, altering the coevolutionary signal to promote alternative predictions [7, 8, 9, 10].

Our work falls within this MSA-manipulation category, where several methodological variants have emerged. Subsampling approaches, such as the work of del Alamo et al. [11], randomly select subsets of sequences to drive structural diversity. Clustering approaches, such as AF-Cluster [12], partition MSAs into representative subsets to induce structural diversity. More recently, purification strategies, such as AF-ClaSeq [13], applies M-fold sampling to classify and “purify” sequence subsets associated with distinct conformational states of dynamic proteins. While these approaches have revealed alternative structures, the combinatorial complexity of MSA space is enormous, making exhaustive exploration infeasible. Consequently, many biologically relevant conformational states likely remain inaccessible to current methods.

To address this limitation, we introduce MSACLR, a contrastive learning framework that aligns MSA space with conformational space. By training on AlphaFold Database (AFDB) Clusters [14] and synthetic datasets, MSACLR learns embeddings that discriminate between structural clusters, enabling efficient exploration of MSA space without invoking any structure module. This provides a foundation for integrating sampling strategies, such as Markov Chain Monte Carlo (MCMC), to recover full conformational ensembles, bridging the gap between static structure predictions and dynamic protein behavior.

## 2 Methods

### 2.1 Stage 1: Pre-Training with AFDB Cluster Dataset

#### 2.1.1 AFDB Cluster Filtering and MSA Construction

We curated protein cluster data from the AFDB Clusters using a systematic multi-criteria filtering pipeline. Clusters were retained only if they satisfied all of the following conditions: (i) cluster size ≥ 10 members; (ii) both the representative structure’s pLDDT score [15] (repPlddt) and average pLDDT score of the protein structures in the Foldseek cluster (avgPlddt) ≥ 90; and (iii) the sequence length of the representative protein (repLen) as well as the average length of the protein sequences in the Foldseek cluster (avgLen) ≤ 500 residues. These thresholds were selected to enrich for compact, well-structured domains while excluding sparsely populated or low-confidence clusters. For each cluster that passed these criteria, we subsampled up to five non-representative members while always retaining the representative structure. This procedure reduced redundancy and maintained balanced coverage across clusters. After filtering and subsampling, the final dataset comprised 33,445 unique clusters and 200,670 unique proteins.

To construct MSAs for these proteins, we employed `MMseqs2-GPU` [16] to search for homologous sequences, yielding 175,722 A3M files. We then applied filtering criteria to ensure quality and computational efficiency: (i) query sequence length ≤ 256 residues; (ii) at least 100 sequences per MSA, and (iii) clusters containing at least two valid members.The final dataset consisted of 24,352 clusters and 118,387 unique proteins, which served as the training data for Stage 1 contrastive learning.

#### 2.1.2 Pre-Training

MSACLR is inspired by the SimCLR framework [17], adapting contrastive learning to MSAs instead of images (Figure 1). The objective is to learn MSA embeddings that align with structural clusters from AFDB (Figure 3b). Let *C*_*n*_ denote the MSA pool corresponding to AFDB cluster *n*. From each cluster *C*_*n*_, we randomly subsample two subMSAs, 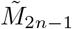 and 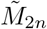, which form a positive pair. SubMSAs drawn from different clusters are treated as negatives.

**Figure 1.**
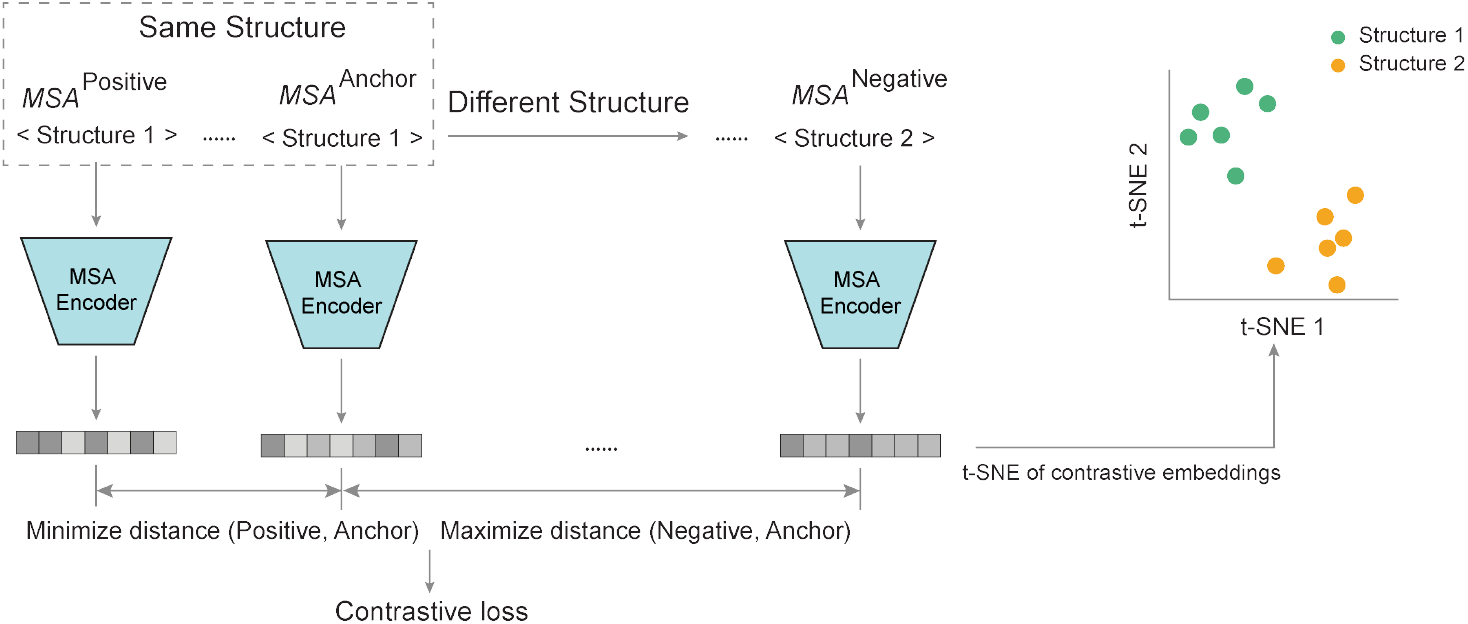
Overview of the MSACLR workflow. MSAs are encoded into embeddings, which are trained with a contrastive loss to minimize the distance between embeddings from the same structural class and maximize the distance between those from different classes. This process yields embeddings that reflect structural relationships among proteins.

Each subMSA is encoded using a LoRA-adapted [18] Evoformer module, producing both MSA representations and pair representations. These representations are pooled and concatenated into a feature vector *h*_*i*_, *h*_*j*_. A non-linear projector *g*(·) maps each feature vector into a 256-dimensional contrastive latent vector *z*_*i*_, *z*_*j*_ (Figure 3a).

The model is trained with the normalized temperature-scaled cross-entropy (NT-Xent) loss [17]:

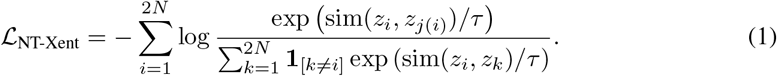

where *z*_*j*(*i*)_ is the positive counterpart of *z*_*i*_, sim(·,·) denotes cosine similarity, and *τ* is the temperature.

### 2.2 Stage 2: Fine-Tuning with Generated SubMSAs Dataset

### 2.3 Generated SubMSAs Dataset

To generate training data for Stage 2 of MSACLR, we selected 3,056 proteins and constructed subMSAs from the full MSA of each protein (Figure 3d). Multiple subMSAs, each containing 8 sequences, were sampled and submitted to `ColabFold` [19] for structure prediction. Predicted structures with pLDDT scores below 70 were discarded to ensure structural quality. The remaining models were clustered using `Foldseek easy-cluster` [20] (with parameters `-c 0.9 --alignment-type 1 --tmscore-threshold 0.85 --exact-tmscore 1`). Based on the clustering results, only proteins with at least 8 structural clusters, each containing at least 4 distinct structural members, were retained. After filtering, this procedure yielded 1,019 proteins.

For supervised contrastive learning, these structural clusters provide labels for their corresponding subMSAs. To further expand the dataset, we applied BLOSUM62-guided substitution to the subMSAs during fine-tuning. Specifically, for each subMSA, 6% of the aligned (non-query) positions were randomly selected, and the original residues were replaced by sampling one of the top-2 candidate substitutions ranked by the BLOSUM62 matrix. Each subMSA was augmented on-the-fly 15 times, ensuring at least 64 subMSAs per structural cluster and a minimum of 8 structural clusters per protein.The augmented subMSAs are treated as positive samples with respect to their original subMSAs.

### 2.4 Fine-Tuning

In Stage 2, MSACLR is fine-tuned to make embeddings discriminative for different predicted structural models of the same protein (Figure 3c). The pretrained MSA encoder is frozen, and only the projector is updated. SubMSAs are labeled according to the clustering of their predicted models, enabling supervised contrastive learning. Within each protein, subMSAs with the same structural label are treated as positives, while those with different labels are treated as negatives.

Let *I* denote the index set of subMSAs in a batch. For each anchor *i* ∈ *I*, let *P* (*i*) denote the set of indices of all other subMSAs in the batch that share the same structural label as *i*. The supervised contrastive loss [21] is defined as:

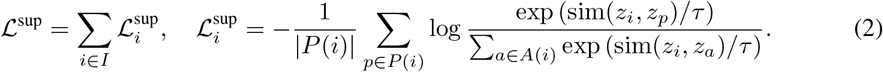

where *z*_*i*_ is the embedding of anchor *i, A*(*i*) denotes all indices in the batch excluding *i*, and *τ* is the temperature.

## 3 Results

### 3.1 First-Stage MSACLR Embeddings Capture Fold-Level Separation

We analyze the embeddings produced by first-stage MSACLR using t-SNE visualization [22], which projects similar embeddings into adjacent points in a two-dimensional space. Figure 4a shows embeddings of proteins from the SCOPe dataset [23] (15,141 proteins), where MSACLR embeddings exhibit clear fold-level separation among the top 10 folds.

To assess whether similar fold separation could be achieved from single-sequence embeddings, we compared MSACLR with the state-of-the-art model ESM-C (6B) [24]. Due to API access limits for ESM-C, this comparison was restricted to proteins from the two largest SCOPe classes, class a (all-*α* proteins) and class b (all-*β* proteins), comprising 5,693 proteins. As shown in Figure 4b, ESM-C embeddings show limited fold-level separation. In contrast, MSACLR embeddings trained on MSAs (Figure 4c) achieve improved clustering, capturing fold distinctions more effectively.

These results highlight two key observations: (1) MSAs provide richer information (stronger signals) than single-sequences for distinguishing structural folds. (2) First-stage MSACLR embeddings reveal improved structural resolution, enabling discrimination at the fold level.

### 3.2 Second-stage MSACLR embeddings capture structural variations within the same protein

Fine-tuned MSACLR embeddings demonstrate the ability to discriminate MSAs associated with different conformations of the same protein. Figure 2 illustrates representative cases, where subMSAs generated by AF-ClaSeq and subsequently labeled through structure prediction and Foldseek clustering form well-separated groups in the embedding space. Additional examples are provided in Figure 5.

**Figure 2.**
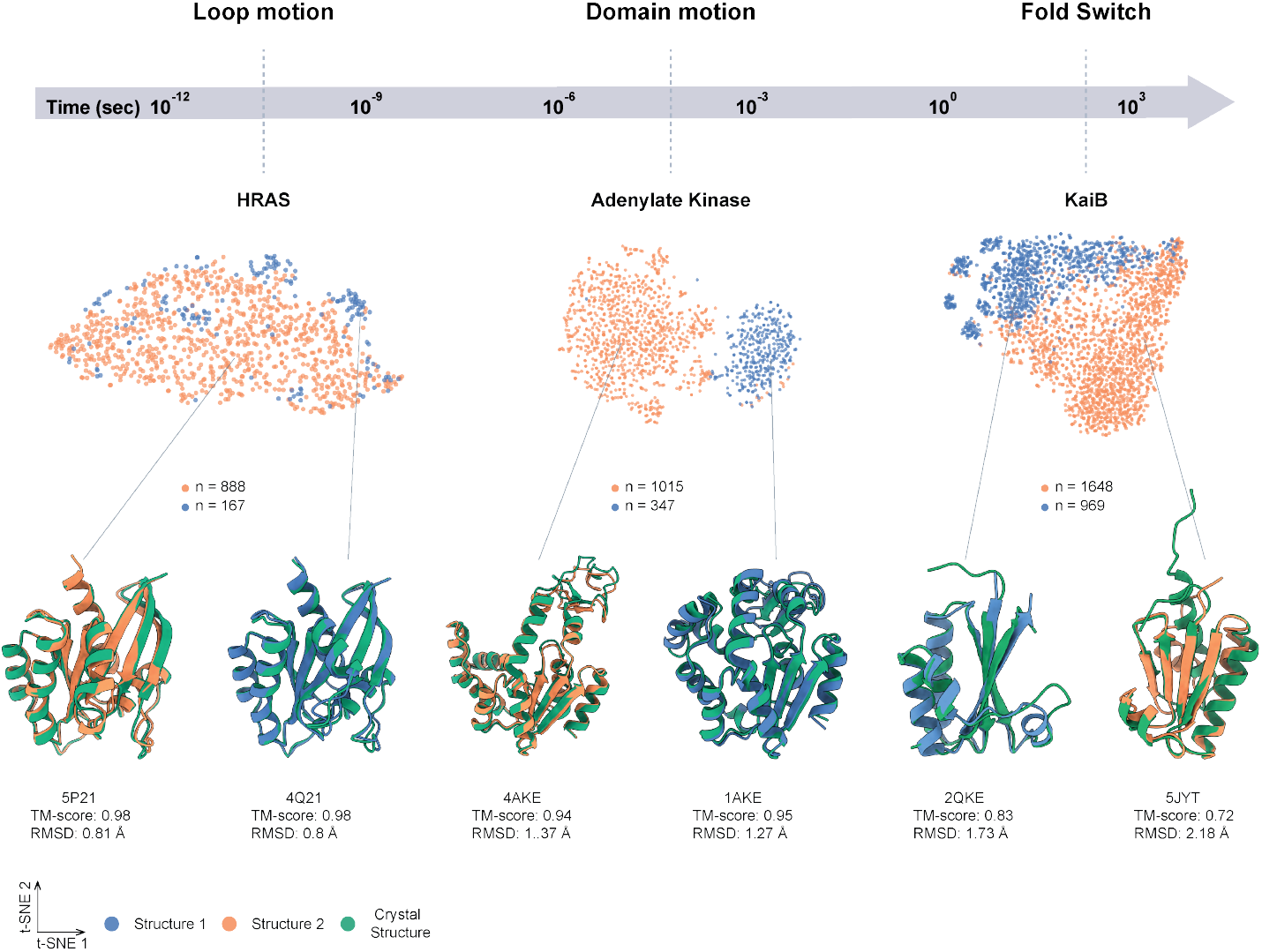
Fine-tuned MSACLR embeddings capture structural differences across timescales. Visualization of embeddings for representative proteins spanning a range of conformational dynamics: local loop rearrangements (HRAS), large domain motions (Adenylate kinase), and fold switching (KaiB). Each point represents a subMSA, and colors indicate state assignments. Experimental structures are shown alongside representative predicted models. Distinct embedding clusters demonstrate that fine-tuned embeddings are sensitive to conformational variation across different timescales.

**Figure 3.**
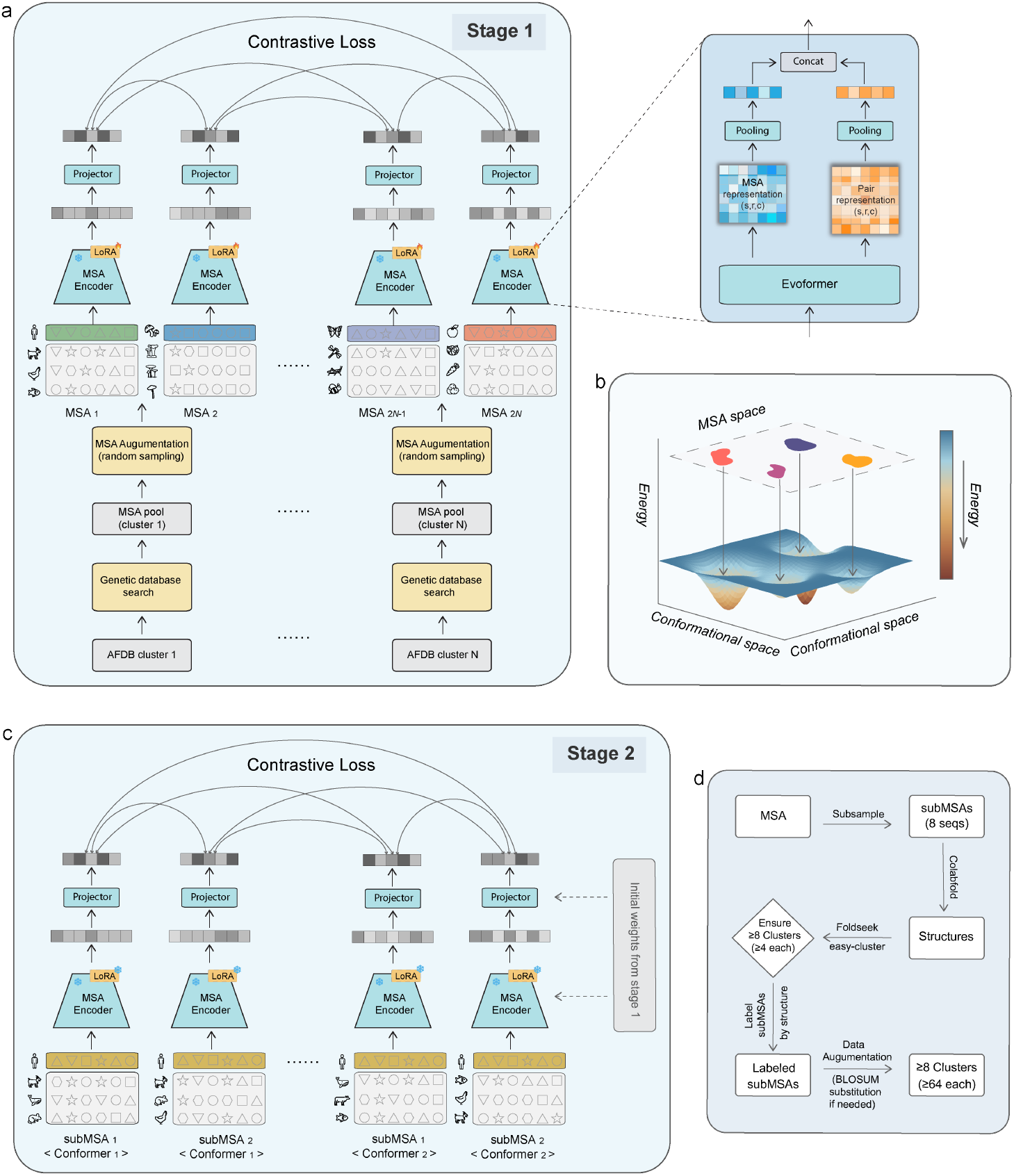
MSACLR: A two-stage contrastive learning framework mapping MSA space to protein conformational space. a, Stage 1(pretraining): MSAs from the same AFDB cluster are treated as positives, while those from different clusters are negatives. Only the LoRA parameters of the MSA encoder are updated, while the base encoder remains frozen. Under a contrastive loss, the LoRA-adapted MSA encoder and projector learn embeddings that capture MSA-structure relationships. In each MSA, the top row (colored) denotes the query sequence, with different colors representing different proteins (queries). b, The goal of MSACLR is to map MSA space into conformational space, where embeddings of MSAs associated with the same conformer cluster together, while those from different conformers remain separated. c, Stage 2 (fine-tuning): SubMSAs corresponding to predicted structure clusters are encoded with weights initialized from Stage 1. Under contrastive loss, the projector is refined to discriminate between MSAs of different structural states for the same protein. In each subMSA, the top row (colored) denotes the query sequence, with identical colors representing the same proteins. d, Data generation workflow: Multiple subMSAs are sampled from the full MSA and submitted to ColabFold for structure prediction. Predicted models are clustered, and each cluster is associated with multiple subMSAs for supervised contrastive learning.

**Figure 4.**
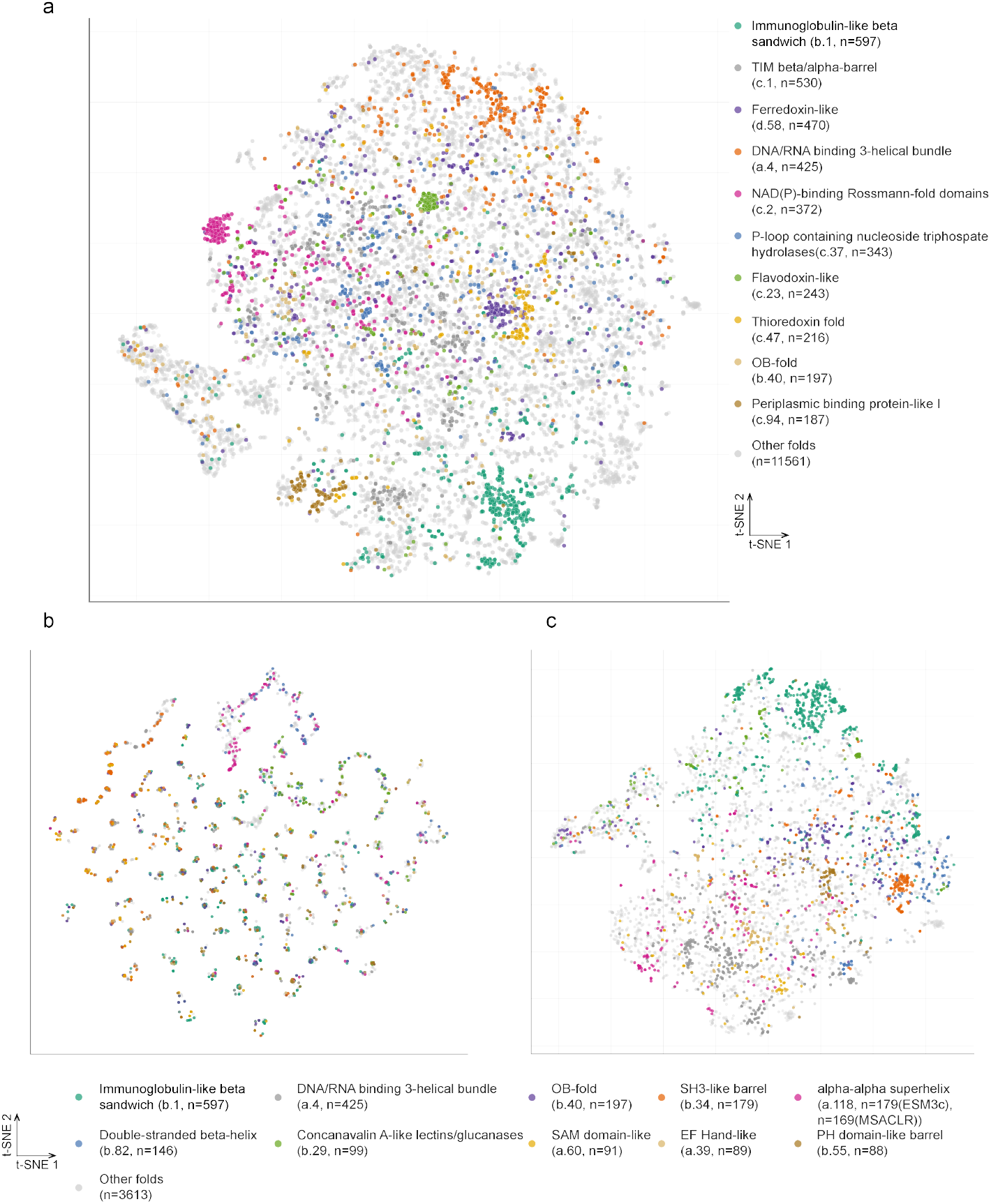
SCOPe dataset Mapping. t-SNE visualizations of protein embeddings showing structural fold distribution. a, Pretrained MSACLR embeddings separate proteins into distinct clusters corresponding to the top 10 folds in the SCOPe dataset. b, Embeddings from the ESM-C (6B) single-sequence model applied to proteins from the top 10 folds within SCOPe classes a and b show poor separation, indicating that single-sequence embeddings fail to capture fold-level information. c, MSACLR embeddings, trained on MSAs, reveal clear clustering of proteins from these top 10 folds, highlighting that MSA-based models capture structural fold information more effectively than single-sequence embeddings.

**Figure 5.**
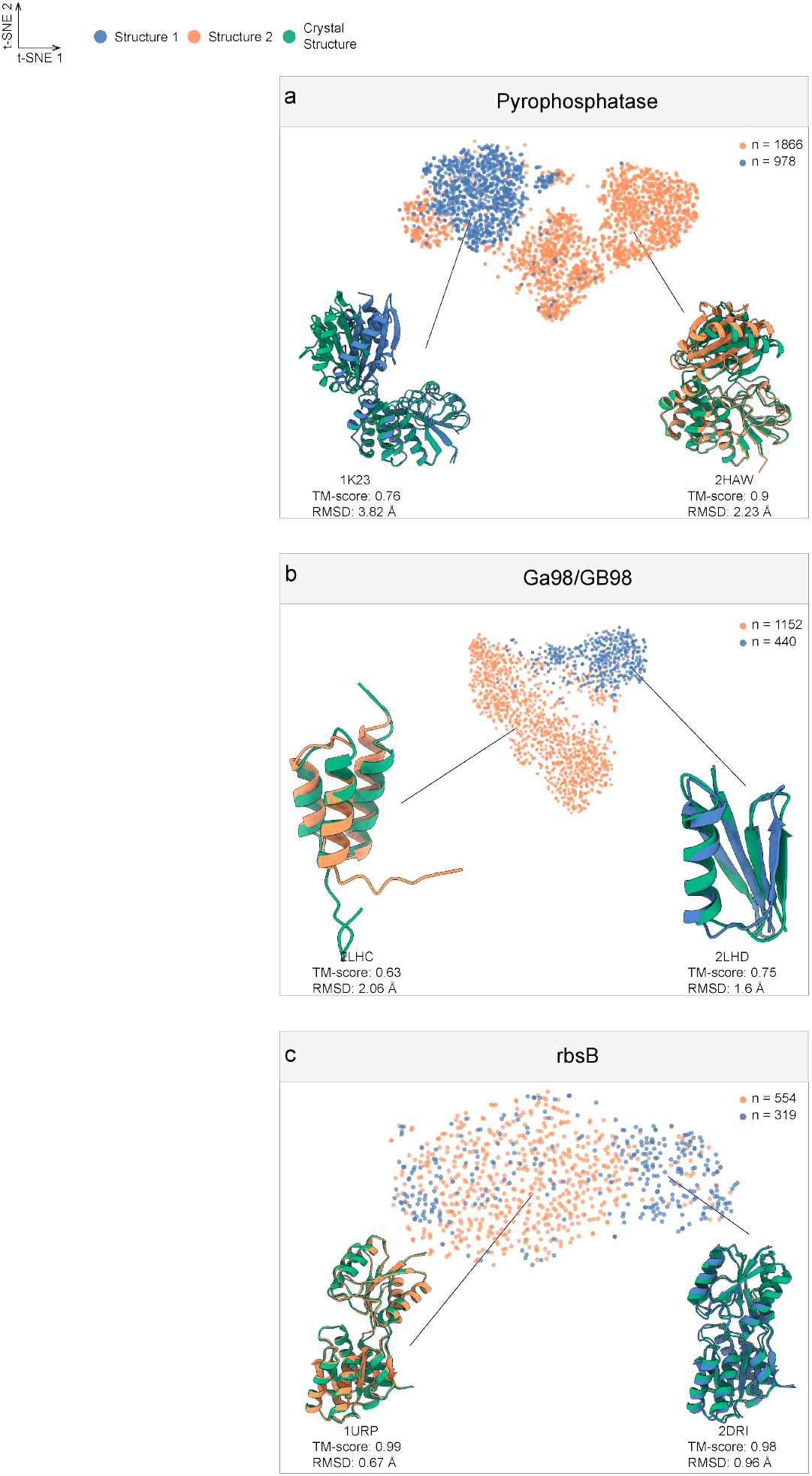
Fine-tuned MSACLR embeddings separate distinct structural states. Visualization of embeddings after fine-tuning, showing that subMSAs corresponding to different predicted structural states of the same protein form distinct clusters in the embedding space. Each point represents a subMSA, and colors indicate state assignments. Experimental structures are shown together with representative predicted structures. Distinct embedding clusters highlight that fine-tuned embeddings capture structural differences among predicted models.

These examples span multiple scales of structural variation. KaiB and GA98/GB98 represent global fold-switch cases; Pyrophosphatase, Adenylate Kinase, and rbsB exhibit large-scale domain motions; while HRAS shows variations confined to local loops. In each case, the fine-tuned MSA embeddings align with their corresponding structural states, suggesting that the model can capture even subtle conformational differences within individual proteins.

These results highlight two key observations: (1) Fine-tuned MSACLR embeddings can associate subMSAs with distinct conformational states of the same protein; and (2) Embeddings are sensitive to structural differences across scales, from local loop motions to large domain motions.

## 4 Conclusion and future work

In this work, we introduced MSACLR, a two-stage contrastive learning framework that maps protein MSA representations to conformational space. Stage 1 captures relationships between MSAs and structural folds across proteins, while Stage 2 targets structural variations within individual proteins. By incorporating BLOSUM62-based augmentation, MSACLR learns representations that reflect how MSA variations relate to structural differences. This mapping establishes a foundation for efficient sequence-space exploration, facilitating multi-state structural prediction and deepen our understanding of protein dynamics.

Future work will explore MSACLR as a surrogate model integrated with sampling methods such as MCMC, enabling efficient navigation of the vast MSA space (Figure 6). Our goal is that, as sampling converges, basins in the MSACLR embedding space will emerge, corresponding to distinct protein conformations. Additionally, MSACLR can be combined with AF-ClaSeq to accelerate the identification of “purified” sequence subsets for predicting alternative conformational states of dynamic proteins while significantly reducing the computational cost of repeated AlphaFold2 predictions.

**Figure 6.**
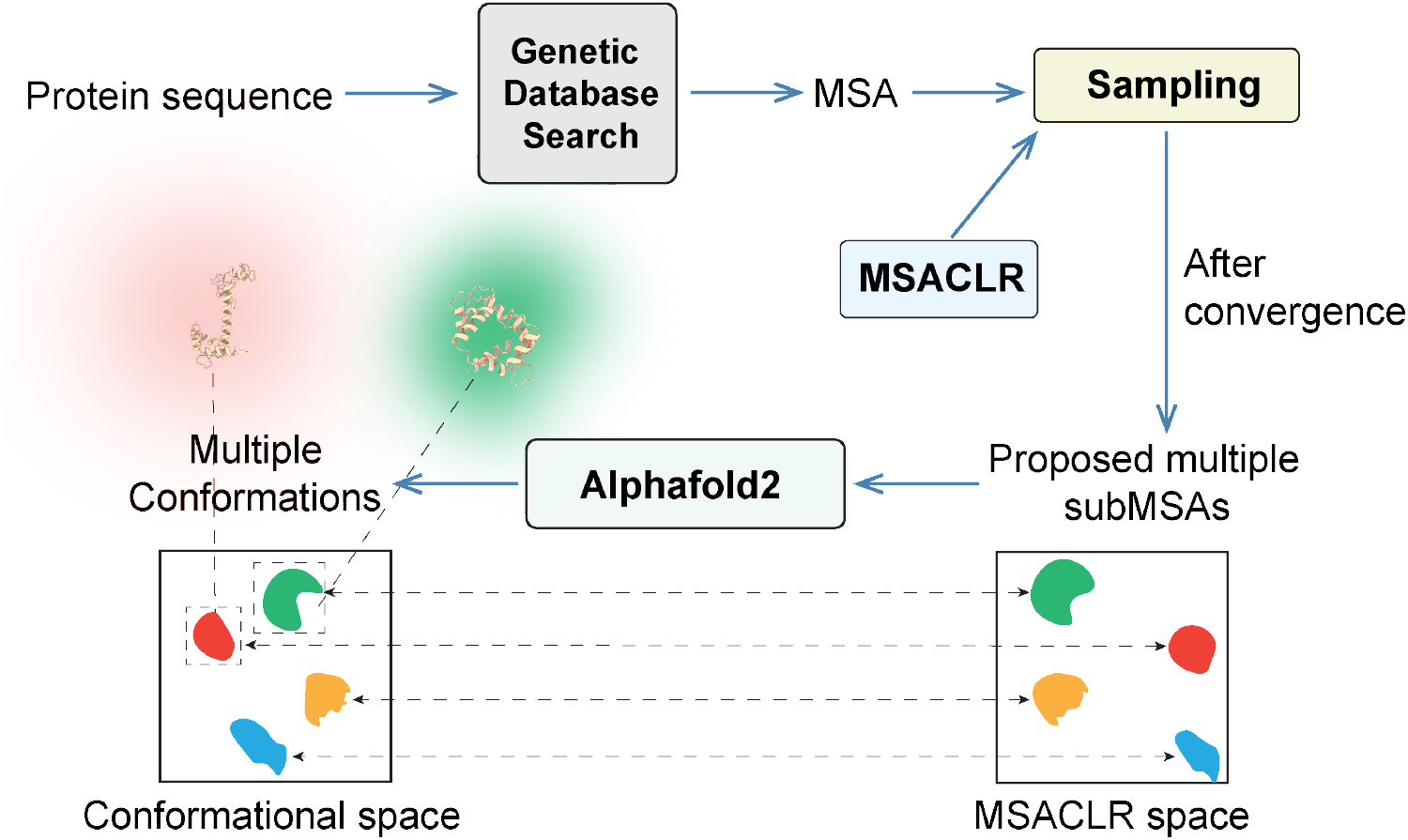
Future extensions will integrate MSACLR with sampling methods to reveal embedding basins corresponding to distinct conformations.

## A Appendix

### A.1 Code Availability

We commit to releasing the source code and trained model weights upon acceptance of this work.

### A.2 Data Availability

We commit to releasing all datasets used in this work upon acceptance of this work.

### A.3 Training Details

#### A.3.1 First-stage MSACLR

Each MSA is encoded using the Evoformer module from the `OpenFold` [25] implementation, with the `max msa clusters` set to 168 and crop size set to 256. The encoded features are passed through an MLP projector with a hidden dimension of 1024, which maps them into a 256-dimensional contrastive latent space. To improve efficiency, we apply LoRA with *r* = 16 and *α* = 32, targeting the Evoformer’s attention modules `(linear z, linear q, linear k, linear v)`.

The model is optimized with the AdamW [26] optimizer using a learning rate of 1.0 × 10^*−*4^ and weight decay 1.0 × 10^*−*4^. A cosine learning rate scheduler with 1,000 warm-up steps is employed over 10,000 total training steps. We train with a batch size of 1,024 under the NT-Xent loss, with temperature *T* = 0.07.

#### A.3.2 Second-stage MSACLR

In Stage 2, the subMSAs sampled from each protein are processed by the Evoformer encoder with `max msa clusters`=10 and `crop size=`268. The encoder is initialized from the pre-trained Stage1 weights and kept frozen during fine-tuning. Only the MLP projector, with a hidden dimension of 1024 mapping into a 256-dimensional contrastive latent space, is updated.

The model is optimized with the AdamW optimizer using a learning rate of 5.0 × 10^*−*5^ and weight decay 1.0 × 10^*−*4^. A cosine learning rate scheduler with 2,500 warm-up steps is employed over 25,000 total training steps. We train with a batch size of 512 under the supervised contrastive loss, with temperature *T* = 0.07.

### A.4 Algorithms

#### Algorithm 1

MSACLR’s main learning algorithm.

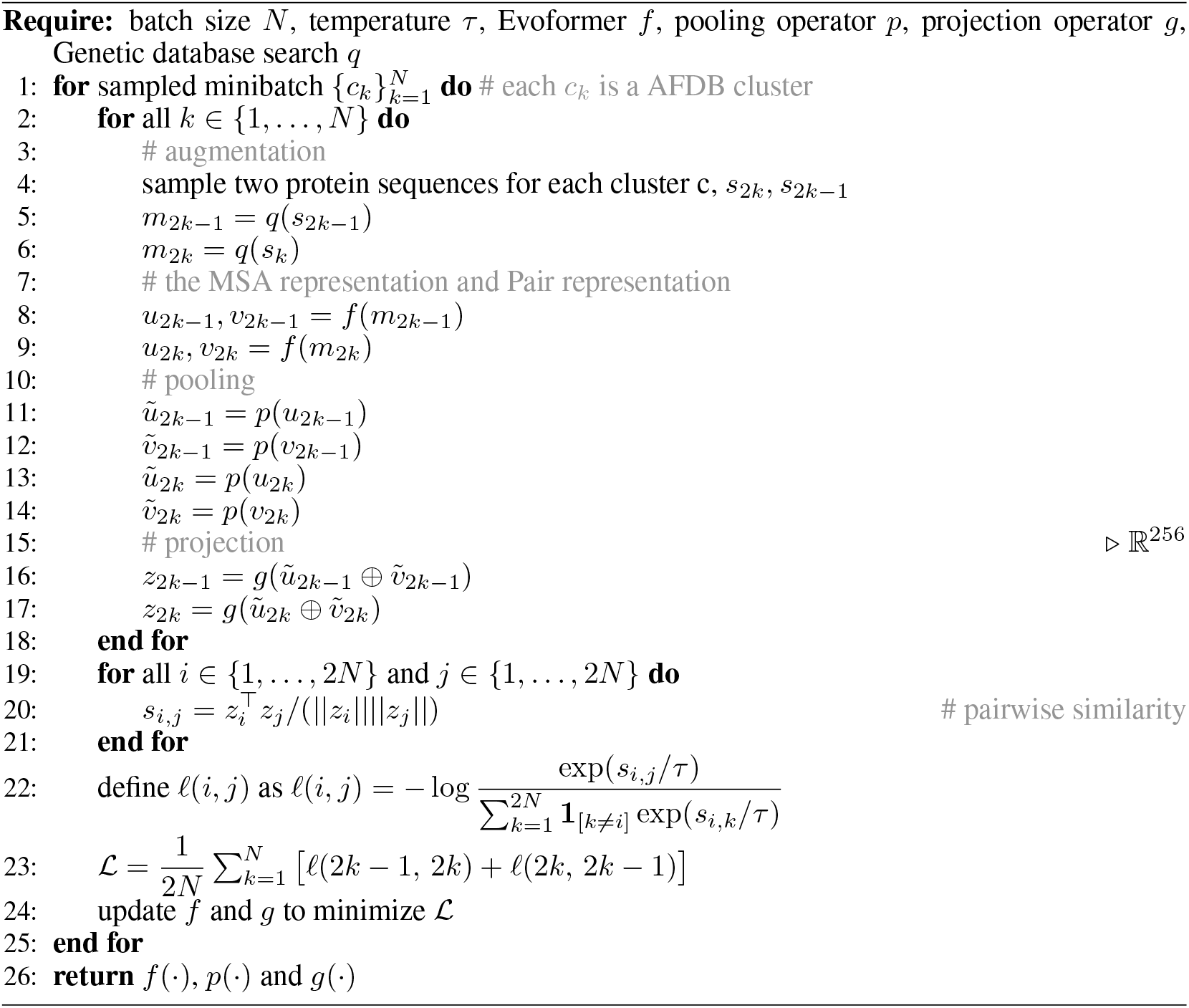

#### Algorithm 2

Projection operator

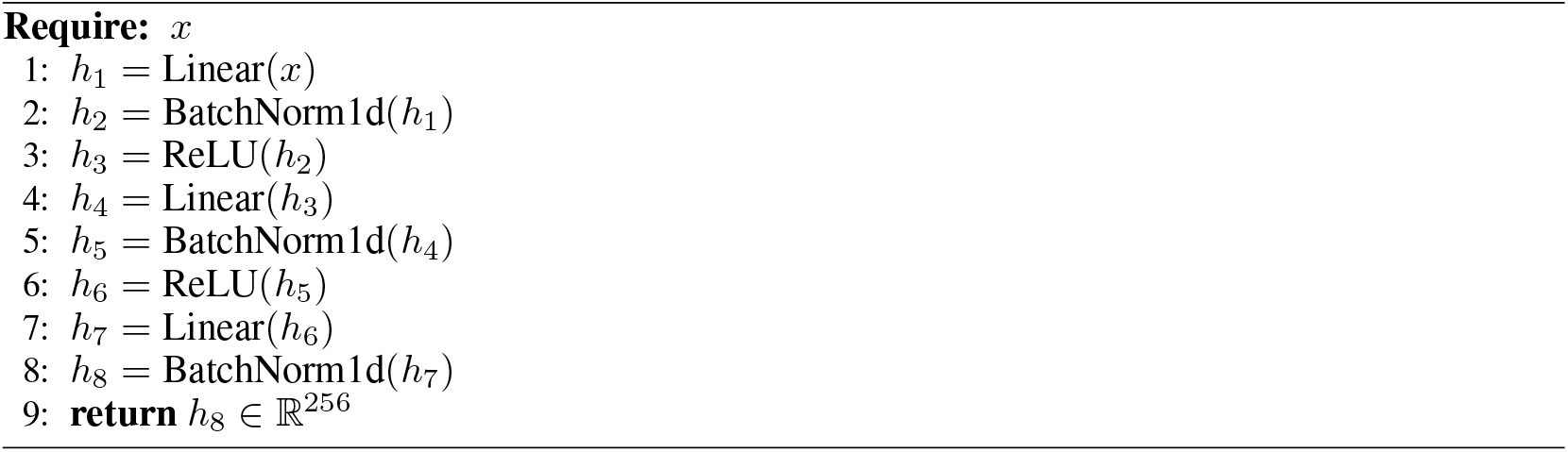

